# *Arabidopsis* histone H3 lysine 9 methyltransferases KYP/SUVH5/6 are involved in leaf development by interacting with AS1-AS2 to repress *KNAT1* and *KNAT2*

**DOI:** 10.1101/2022.02.23.481647

**Authors:** Fu-Yu Hung, Yun-Ru Feng, Yuan-Hsin Shih, You-Cheng Lai, Keqiang Wu

## Abstract

*Arabidopsis* KRYPTONITE/SUPPRESSOR OF VARIEGATION 3–9 HOMOLOG 4 (KYP/SUVH4), SUVH5 and SUVH6 are H3K9 methyltransferases and they are redundantly involved in silencing of transposable elements (TEs). A recent study indicated that KYP/SUVH5/6 can directly interact with the histone deacetylase HDA6 to synergistically regulate TE expression. However, the function of KYP/SUVH5/6 in plant development is still unclear. The ASYMMETRIC LEAVES1 (AS1) and AS2 form a transcription complex, which is involved in leaf development by repressing the homeobox genes *KNOTTED-LIKE FROM ARABIDOPSIS THALIANA 1* (*KNAT1*) and *KNAT2*. In this study, we found that KYP and SUVH5/6 directly interact with AS1-AS2 to repress *KNAT1* and *KNAT2* by altering histone H3 acetylation and H3K9 dimethylation levels. In addition, KYP can directly target on the promoters of *KNAT1* and *KNAT2*, and the binding of KYP is dependent on AS1. Furthermore, the genome-wide occupancy profile of KYP indicated that KYP is enriched in the promoter regions of coding genes, and the binding of KYP is positively correlated with that of AS1 and HDA6. Together, these results indicate that *Arabidopsis* H3K9 methyltransferases KYP/SUVH5/6 are involved in leaf development by interacting with AS1-AS2 to alter histone H3 acetylation and H3K9 dimethylation from the *KNAT1* and *KNAT2* loci.

## Introduction

The initiation of leaf primordia is established by recruitment of the cells flanking the shoot apical meristem (SAM). Meristem activity in the shoot apex is specified in part by the class I *KNOTTED-LIKE HOMOBOX* (*KNOX*) genes (Long et al., 1996; Vollbrecht et al., 2000; Scofield and Murray, 2006). Lateral organs, such as leaves, initiate on the flank of the shoot apical meristem, and down-regulation of *KNOX* gene expression is essential to facilitate this process (Jackson et al., 1994; Long et al., 1996). Moreover, silencing of *KNOX* genes is important in developing organs since ectopic *KNOX* expression during organogenesis results in patterning defects and hyper-proliferation of cells (Sinha et al., 1993; Chuck et al., 1996; Kidner et al., 2002). In *Arabidopsis*, the members of the *KNOX* family can be divided into three classes. Class I KNOX genes include *BREVIPEDICELLUS/KNOTTED-LIKE FROM ARABIDOPSIS THALIANA1* (*BP/KNAT1*), *KNAT2*, *KNTA6* and *SHOOTMERISTEMLESS* (*STM*) (Byrne, 2005). Class II *KNOX* genes comprise *KNAT3*, *KNAT4*, *KNAT5* and *KNAT7*, which are broadly expressed and have been shown to function redundantly to influence lateral organ differentiation in *Arabidopsis* (Furumizu et al., 2015). Class III only contains *KNATM*, which is a novel *KNOX* gene lacking the homeodomain (Magnani and Hake, 2008). In *Arabidopsis*, *KNAT1* is expressed in the vegetative meristem and stem, and is down-regulated as leaf primordia are initiated (Chuck et al., 1996). Thus, the precise balance between the differentiation and proliferation of stem cells is achieved in part through proper regulation of *KNOX* expression.

*KNOX* repression during organogenesis is mediated by the transcription complex composed of the MYB domain protein ASYMMETRIC LEAVES1 (AS1) and the LOB domain protein AS2 in *Arabidopsis* (Timmermans et al., 1999; Tsiantis et al., 1999; Byrne et al., 2000; Ori et al., 2000; Iwakawa et al., 2002). *KNAT1* and *KNAT2* are mis-expressed in the leaves and flowers of the *as1/as2* double mutant, suggesting that AS1 and AS2 promote leaf differentiation through repressing *KNOX* (Ori et al., 2000). The AS1-AS2 complex (AS1/2) can recruit a chromatin-remodeling protein HISTONE REGULATORY HOMOLOG 1 (HIRA) to regulate target gene expression during organogenesis (Guo et al., 2008). In addition, AS1/2 can also recruit POLYCOMB-REPRESSIVE COMPLEX 2 (PRC2) to repress *KNOX* genes by histone H3 lysine 27 methylation (Lodha et al., 2013). Collectively, these studies suggested that the repression activity of AS1/2 is associated with histone modifications.

Histone modifications including methylation, acetylation, phosphorylation and ubiquitination can influence transcription, DNA repair, replication and recombination (Klose and Zhang, 2007; Berger, 2007). Lysine methylation on the side chains of histones is regulated by histone methyltransferases (HMTs) and histone demethylases (HDMs) (Klose and Zhang, 2007; Berger, 2007). Methylation on lysine 9 and 27 of histone H3 (H3K9me and H3K27me) is associated with transcription repression, while methylation on lysine 4 and 36 of histone H3 (H3K4me and H3K36me) is associated with transcription activation (Klose and Zhang, 2007; Berger, 2007). For instance, H3K9 mono-methylation (H3K9me1) and H3K9 dimethylation (H3K9me2) mainly function in repressing transposon activities. H3K9me2 is enriched in transposons and repeated sequences (Liu et al., 2010; Bernatavichute et al., 2008; Stroud et al., 2014; Du et al., 2015). In addition, the level of histone acetylation is controlled by histone acetyltransferases (HATs) and histone deacetylases (HDACs). HATs can add acetyl groups on lysine, which loosens chromatin confirmation and leads to transcription activation. On the contrary, removing acetyl groups from lysine by HDACs leads to condensed chromatin structure and transcription repression (Klose and Zhang, 2007; Berger, 2007).

Histone lysine methyltransferases (HKMTs) have a specific conserved domain called SET (SUPPRESSOR OF VARIEGATION, ENHANCER OF ZESTE AND TRITHORAX) domain, which is mainly responsible for histone methylation activity. In *Arabidopsis*, 49 SET Domain Group (SDG) proteins have been identified, and 31 of them are known or predicted to have HKMT activity. These SDG proteins can be further classified into five classes (class I to V) based on their domain architectures or their target lysine residues (Ng et al., 2007). Previous studies have revealed that the Class V SDG proteins including SUPPRESSOR OF VAR-IEGATION 3–9 HOMOLOG (SUVH) and SUPPRESSOR OF VARIEGATION 3–9 RELATED (SUVR) proteins are associated with H3K9 methylation involved in heterochromatin maintenance and DNA methylation (Grafi et al., 2007; Johnson et al., 2007; Pontes et al., 2006; Pontvianne et al., 2010). All SUVH proteins contain a SET domain, a pre-SET domain, a post-SET domain, and a STE and RING-associated (SRA) domain. The SRA domain is responsible for recognizing methylated DNA (Du et al., 2014). KRYPTONITE (KYP, also called SUVH4), SUVH5 and SUVH6 are the best characterized SUVH proteins in *Arabidopsis* and they function as histone H3K9 methyltransferases. KYP is required for the maintenance of CHG methylation controlled by CHROMOMETHYLASE 3 (CMT3) (Jackson et al., 2002; Tran et al., 2005; Ebbs and Bender, 2006). Furthermore, KYP, SUVH5 and SUVH6 act redundantly to silence transposable elements (TEs) by regulating H3K9me1 and H3K9me2 at their target loci. The *kyp/suvh5/suvh6* triple mutant displays a loss of non-CG methylation similar to the *cmt3* mutant (Tran et al., 2005; Ebbs and Bender, 2006; Johnson et al., 2007; Bernatavichute et al., 2008; Du et al., 2012; Zemach et al., 2013; Stroud et al., 2014). The histone deacetylase HDA6 is also involved in transposon silencing (Liu et al., 2012; Yang et al., 2020). In addition, HDA6 interacts and functions synergistically with KYP, SUVH5 and SUVH6 to co-regulate transposon silencing through histone H3K9 methylation and H3 deacetylation (Yu et al., 2017).

Although it has been established that KYP/SUVH5/6 are important regulators of *TE* silencing, their function in plant development remains elusive. In this study, we found that KYP/SUVH5/6 regulate leaf development by interacting with AS1/2 to repress *KNAT1* and *KNAT2* expression through H3K9me2 and H3 deacetylation.

## Results

### *Arabidopsis* KYP/SUVH5/6 are involved in leaf development

Although previous studies have revealed that several *Arabidopsis* H3K9 demethylases are associated with plant developmental processes (Saze et al., 2008; Dutta et al., 2017; Hung et al., 2020a; Hung et al., 2021), the function of KYP and SUVH5/6 in plant development remains elusive. Our recent study has revealed that KYP and SUVH5/6 interact with the histone deacetylase HDA6 and they function synergistically to regulate TE expression (Yu et al., 2017). To further investigate the biological function of KYP/SUVH5/6, we analyzed the growth phenotypes of *hda6* and *kyp* single, *kyp/hda6* double, *kyp/suvh5/6* triple, and *hda6/kyp/suvh5/6* quadruple mutants. As reported in our previous study (Luo et al., 2012), the *hda6* mutant had curling and serrated leaves. Compared to Col-0 wild type (WT), *hda6, kyp* and *kyp/suvh5/6* mutants displayed a slight curling leaf phenotype (Fig. 1A-D). The curling leaf phenotype was enhanced in *kyp/hda6*, and greatly enhanced in *hda6/kyp/suvh5/6* quadruple mutants. Furthermore, the leaves of *hda6/kyp/suvh5/6* plants were also much smaller (Fig. 1A-D). Quantitative analyses indicated that nearly 80% of leaves in the *hda6/kyp/suvh5/6* quadruple mutant were developmental defected (Fig. 1E, Fig. S1). These results suggest that HDA6 may function synergistically with KYP/SUVH5/6 in the regulation of leaf development.

**Figure 1.**
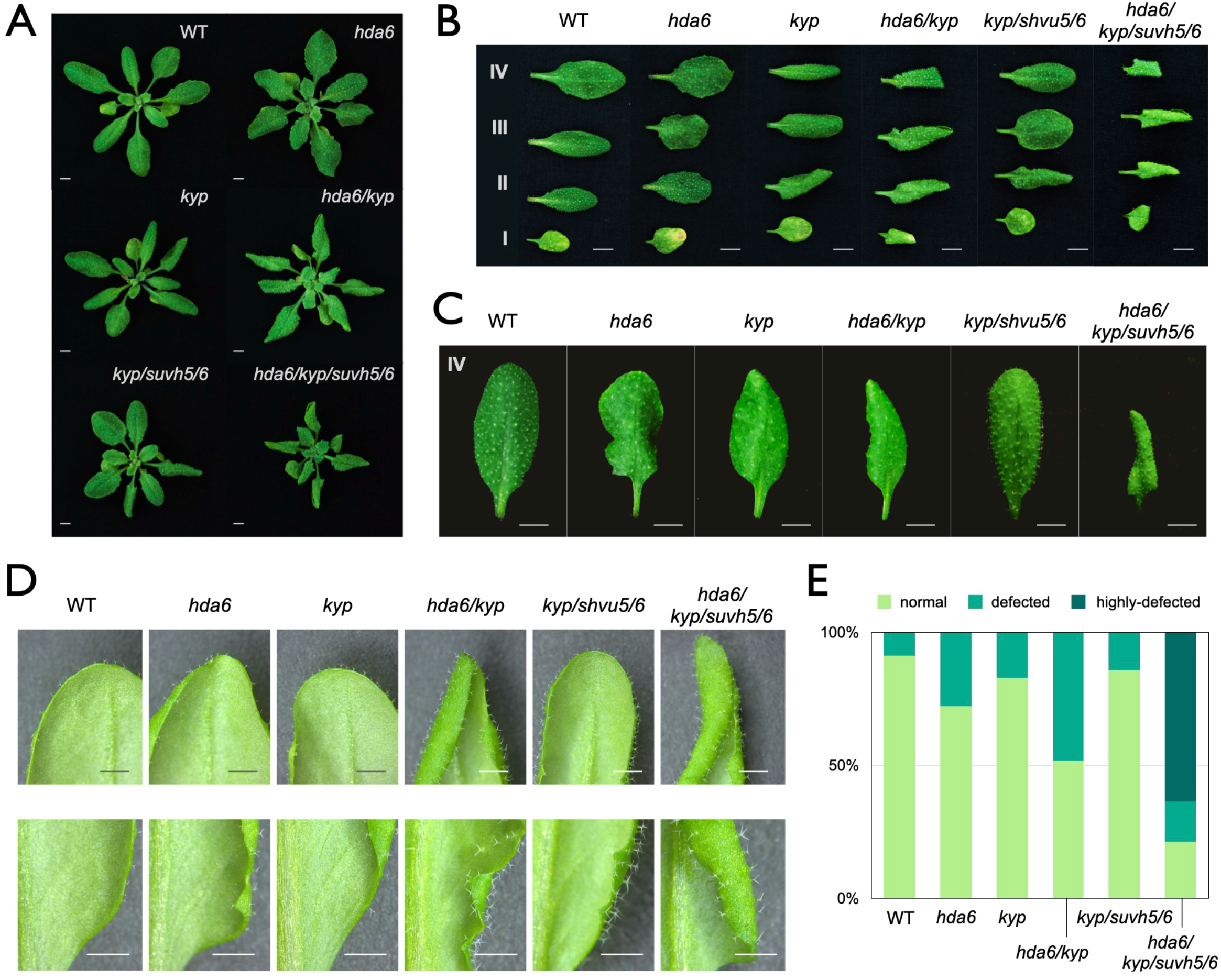
*Arabidopsis* KYP, SUVH5 and SUVH6 are involved in leaf development. (A-D) Leaf development phenotype of WT, *hda6*, *kyp*, *kyp/hda6*, *kyp/suvh5/6* and *hda6/kyp/suvh5/6* mutant plants. The plants were grown under long-day for 20 days. I, II, III and IV indicate the first, second, third and fourth pair of leaf, respectively. Bars= 5mm(A-C) or 2mm(D). (E) Quantitative analysis of leaf development phenotypes of WT, *hda6*, *kyp*, *kyp/hda6*, *kyp/suvh5/6* and *hda6/kyp/suvh5/6* mutant plants. The fourth pair rosette leaves were classified as normal, defected and highly-defected leaves. Frequencies are calculated by the ratio of defected and total leaves examined. At least 40 leaves for each line were scored.

### KYP/SUVH5/6 interact with AS1/2

Our previous study showed that *Arabidopsis* HDA6 Is functionally associated with AS1/2 (Luo et al., 2012). To further investigate the function of KYP and SUVH5/6 in leaf development, we performed bimolecular fluorescence complementation (BiFC) assays and co-immunoprecipitation (Co-IP) assays to investigate whether KYP/SUVH5/6 can interact with AS1/2. We found that KYP, SUVH5 and SUVH6 can interact with both AS1 and AS2 in BiFC assays by using *Agrobacterium*-infiltrated tobacco leaves (Fig. 2A-C) and *Arabidopsis* protoplasts (Fig. S2). The interaction of KYP with AS1 was further confirmed by Co-IP assays using *KYPpro::KYP:GFP*/*kyp* transgenic plants, which carrying KYP fused with GFP driven by the *KYP* native promoter. *35Spro::GFP* transgenic plants carrying GFP driven by the CaMV 35S promoter were used as a control, and the endogenous AS1 protein was detected by using an anti-AS1 antibody. As shown in Figure 2D, AS1 interacted with KYP in Co-IP assays.

**Figure 2.**
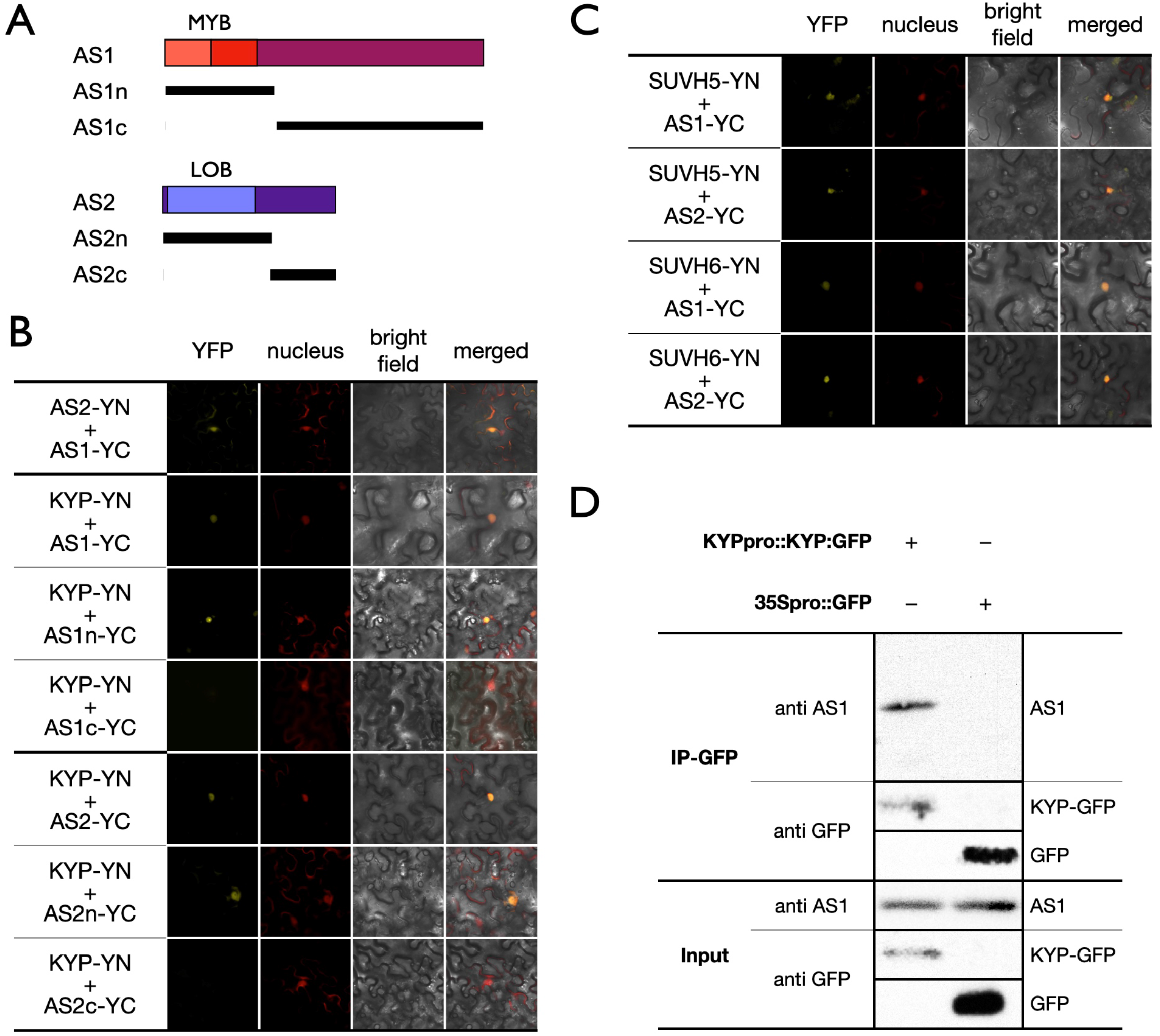
KYP, SUVH5 and SUVH6 interact with AS1 and AS2. (A) Schematic representation of deletions in AS1 and AS2 constructs. MYB: MYB domain of AS1; LOB: LOB-domain of AS2. (B, C) BiFC assays in *N. benthamiana* leaves showing interaction of KYP (B), SUVH5 and SUVH6 (C) with AS1 and AS2 in living cells. Full length KYP, SUVH5, SUVH6 and different regions of AS1/2 were fused with the N terminus (YN) or C terminus (YC) of YFP and co-delivered into tobacco leaves by *Agrobacterium* GV3101. The nucleus was indicated by mCherry carrying a nuclear localization signal. (D) Co-IP of the *KYP* native promoter driven *KYP:GFP* (*KYPpro::KYP:GFP*) or *35Spro::GFP in* transformed *Arabidopsis*. Western blot (WB) was performed with the indicated antibodies.

Various deletion constructs of AS1 and AS2 were also generated to determine the domains responsible for their interaction with KYP in BiFC assays (Fig. 2A). Although the interaction was significantly decreased between KYP and the C-terminus of AS1, the N-terminus of AS1 could still strongly interact with KYP (Fig. 2B). Similarly, the YFP signal could be detected in the nucleus when KYP co-expressed with the N-terminus of AS2, but not with the C-terminus of AS2 (Fig. 2B). These data indicate that the N-terminus of AS1 or AS2 is responsible for the interaction.

In addition, the leaf development phenotype of the *hda6/kyp/suvh5/6* quadruple mutant displayed a defected leaf phenotype similar to *as1* and *as2* (Fig. S3A). We also generated the *as1/kyp* double*, as1/hda6/kyp* triple and *as1/hda6/kyp/suvh6* quadruple mutant plants. Compared to WT, these mutants also displayed a defected leaf phenotype similar to the *as1* (Fig. S3B), suggesting that the function of KYP/SUVH5/6-HDA6 in leaf development is at least partially depending on AS1. Collectively, these results indicated that KYP, SUVH5 and SUVH6 may be involved in leaf development by interacting with AS1 and AS2.

### KYP/SUVH5/6 repress *KNAT1/2* by altering H3K9me2 and H3Ac levels of the *KNAT1/2 loci*

AS1 and AS2 are transcription repressors of the class I *KNOX* genes (Guo *et al*. 2008). To investigate whether KYP and SUVH5/6 affect the expression of *KNOX* genes, we analyzed expression of *KNAT1*, *KNAT2*, and *STM* in WT, *hda6*, *kyp*, *hda6/kyp*, *kyp/suvh5/6* and *hda6/kyp/suvh5/6*. The expression of *KNAT1* and *KNAT2* was significantly increased in the mutants compared to WT (Fig. 3A). Furthermore, the highest expression levels of these class I *KNOX* genes were observed in the *hda6/kyp/suvh5/6* quadruple mutant (Fig. 3A), indicating that KYP, SUVH5/6 and HDA6 can act synergistically to repress the expression of the class I *KNOX* genes.

**Figure 3.**
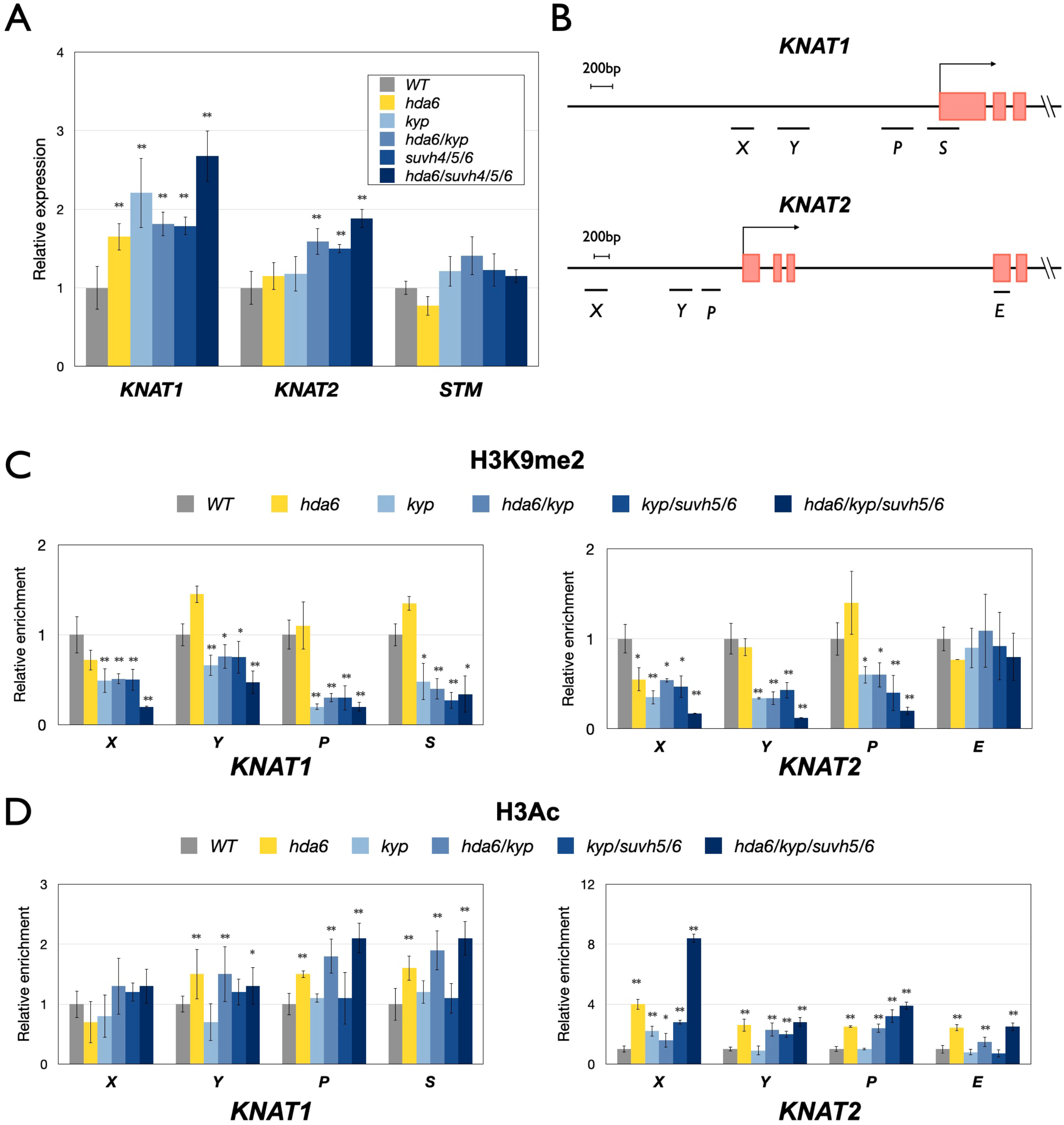
KYP, SUVH5, SUVH6 and HDA6 synergistically regulate *KNAT1* and *KNAT2* by H3K9me2 and H3 deacetylation. (A) Expression of *KNOX* genes in *hda6*, *kyp*, *had6/kyp*, *kyp/suvh5/6*, and *hda6/suvh4/5/6* mutants. `Gene` expression was analyzed by qRT-PCR. RNA was extracted from 10-day-old plants grown under LD conditions. *UBQ10* was used as an internal control. Error bars indicate SD. **P < 0.01 by t-test. At least three independent biological replicates were performed with similar results. (B) Schematic diagram of *KNAT1* and *KNAT2* genomic sections. X, Y and P: promoter regions, S: first exon, E: coding region. (C, D) ChIP analysis of H3K9me2(C) and H3Ac(D) levels on *KNAT1* and *KNAT2* in 10-day-old plants. The amounts of DNA after ChIP were quantified by qPCR and normalized to control. Data points represent average of three technical replicates. Error bars correspond to standard deviations from three biological replicates. *P< 0.05, **P< 0.005 (Student’s t-test).

We further investigated whether KYP/SUVH5/6 and HDA6 affect the level of H3K9me2 and H3Ac on the *KNAT1* and *KNAT2* loci by chromatin immunoprecipitation followed by quantitative PCR (ChIP-qPCR). Several genomic regions of *KNAT1* and *KNAT2* including the AS1/2 binding regions were selected for ChIP-qPCR analysis (Fig. 3B). Compared to WT, we found that the H3K9me2 level of *KNAT1* and *KNAT2* was decreased in *kyp*, *hda6/kyp*, *kyp/suvh5/6* and *hda6/kyp/suvh5/6*, but not in *hda6* (Fig. 3C). In addition, the H3Ac level of *KNAT1* and *KNAT2* was increased in *hda6* and *hda6/kyp/suvh5/6* compared to WT (Fig. 3C). These results suggest that KYP/SUVH5/6 and HDA6 regulate *KNAT1/2* expression through H3K9me2 and H3 deacetylation. Interestingly, we found that the H3K9me2 level on *KNAT1/2* was not decreased in *hda6* (Fig. 3C). Furthermore, the H3Ac level of *KNAT1/2* was less affected in *kyp* and *kyp/suvh5/6* than in *hda6* (Fig. 3C). The expression of *KNAT1/2* was highest in the *hda6/kyp/suvh5/6* quadruple mutant (Fig. 3A), indicating that both decreased H3K9me2 and increased H3Ac contribute to *KNAT1/2* expression changes.

The ChIP assays were used to identify whether KYP can directly target to the *KNAT1* and *KNAT2. KYPpro::KYP:FLAG/kyp* transgenic lines in which the *KYP* genome sequence containing its native promoter fused with the 3xFLAG epitope tag was transformed into the *kyp* background. The *KYP* transcript and protein were detected in the *KYP-pro::KYP:FLAG/kyp* transgenic lines (Fig. S4). ChIP assays were performed with the anti-FLAG antibody using *KYPpro::KYP:FLAG* transgenic seedlings and the binding of KYP to *KNAT1* and *KNAT2* was analyzed by ChIP-qPCR. We found that KYP was highly enriched in the promoter regions of *KNAT1* and *KNAT2* (Fig. 4A, 4B). Furthermore, these KYP-enriched promoter regions are highly overlapping with the binding regions of AS1/2 (Guo et al. 2008). These results indicate that KYP regulates *KNAT1* and *KNAT2* expression by directly targeting on the *KNAT1* and *KNAT2* promoters.

**Figure 4.**
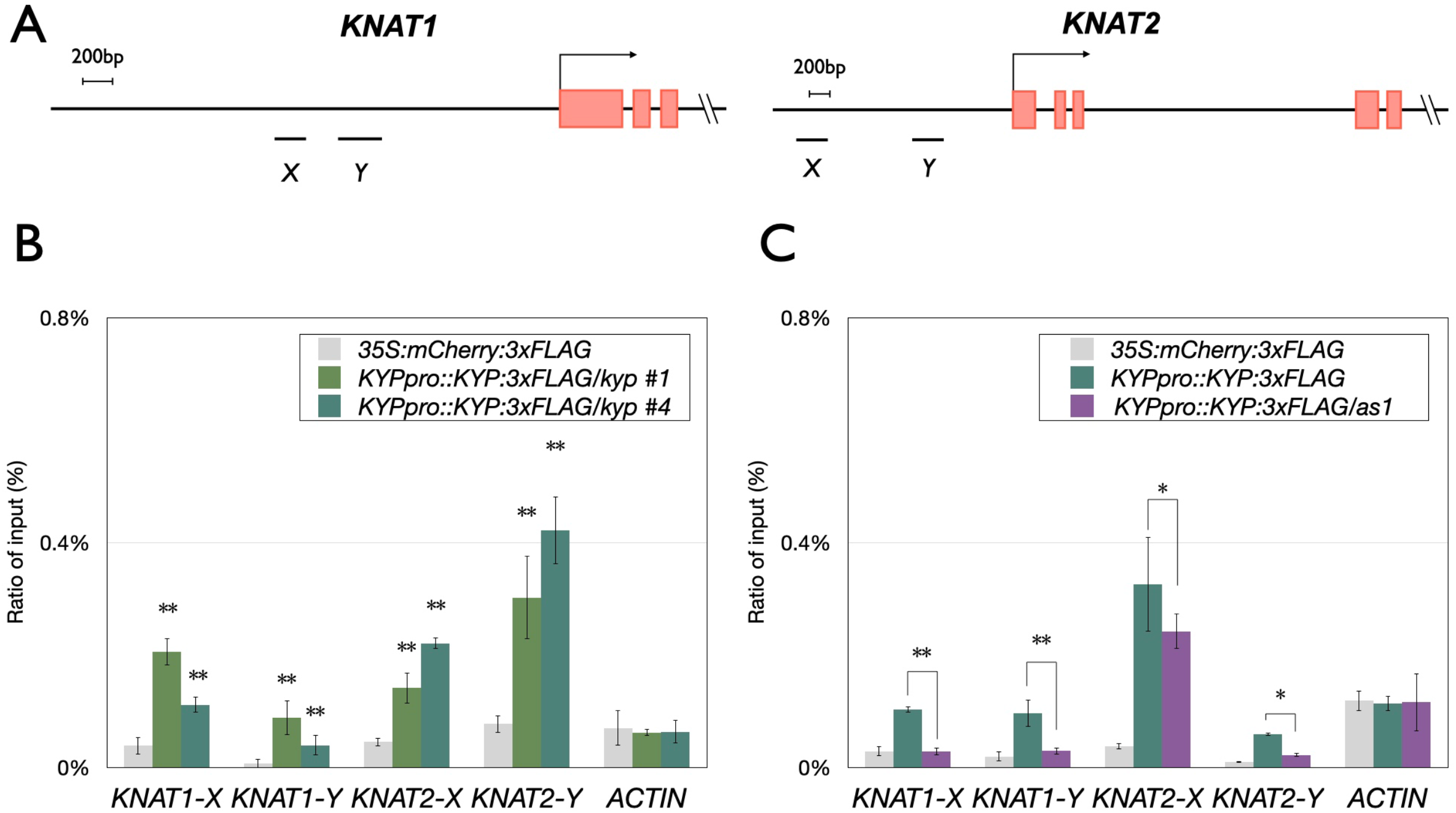
Binding of KYP to *KNAT1* and *KNAT2 in vivo*. (A) Schematic diagram of *KNAT1* and *KNAT2* genomic sections. X and Y regions are the binding sites of AS1/2. (B, C) KYP binding to *KNAT1* and *KNAT2* promoters. 10 days-old seedlings grown under LD were harvested. ChIP assays were performed with the anti-FLAG antibody. The amount of immunoprecipitated DNA was quantified by qPCR. Values represent the normalized average immunoprecipitation efficiencies (%) against the total input DNA. Error bars correspond to standard deviations from three biological replicates. *: P< 0.05, **: P< 0.005 (Student’s t-test).

To further identify the functional correlation between KYP and AS1, we expressed *KYP-pro::KYP:FLAG* in the *as1* mutant background (*KYPpro::KYP:FLAG/as1*). The protein level of KYP in *KYPpro::KYP:FLAG/as1* was similar to that of *KYPpro::KYP:FLAG* (Fig. S4C). We found that the binding of KYP to *KNAT1* and *KNAT2* was significantly reduced in *KYP-pro::KYP:FLAG/as1* (Fig. 4C), indicating that the binding of KYP to the *KNAT1* and *KNAT2* is at least partially dependent on AS1. Furthermore, we also identified the decreased H3K9me2 and increased H3Ac levels on *KNAT1* and *KNAT2* loci in *as1/as2* (Fig. S5), suggesting that AS1/2-regulated *KNAT1* and *KNAT2* expression is associated with H3K9me2 demethylation and H3 acetylation.

### Genome-wide occupancy profiles of KYP

To investigate the genome wide function of KYP in gene regulation, we mapped the genome-wide occupancy of KYP by chromatin immunoprecipitation followed by sequencing (ChIP-seq) using the *KYPpro::KYP:3xFLAG/kyp* transgenic line. 3,924 KYP-targeted genomic regions including *KNAT1* and *KNAT2* were identified. The genome browser views of the ChIP-Seq data show that KYP can target on the *KNAT1* and *KNAT2* promoter regions (Fig. 5A), which is consistent with our ChIP qPCR data. Compared to the *Arabidopsis* genomic region distribution, we found that the binding of KYP was more enriched in the 1kb promoter regions, but less enriched in the gene bodies (Fig. 5B). We also found that most of the KYP targeted genes are protein coding genes (Fig. 5C). We further compared the binding profile of KYP among the protein coding genes and TE genes. In both protein coding genes and TE genes, the general binding of KYP was more enriched on the promoter but less enriched on the gene body (Fig. 5C). Furthermore, we found that the binding of KYP is strongly enriched near the upstream regions of transcription start sites (TSS) of protein coding genes, but not in TE genes (Fig. 5C). These results support that the function of KYP is associated with the regulation of protein coding genes. The KYP targeted genes were further analyzed according to Gene Ontology Biological Processes (GO-BP). We found that the KYP targeted genes are involved in different types of abscisic acid (ABA)/stress responses and development pathways (Fig. 5E). These results are consistent with the finding that KYP is involved in ABA and stress responses (Zheng et al., 2012). Interestingly, we also found that the GO term “leaf development (GO:0048366)” was enriched in the KYP targeted genes (Fig. 5E),supporting a role for KYP in leaf development.

**Figure 5.**
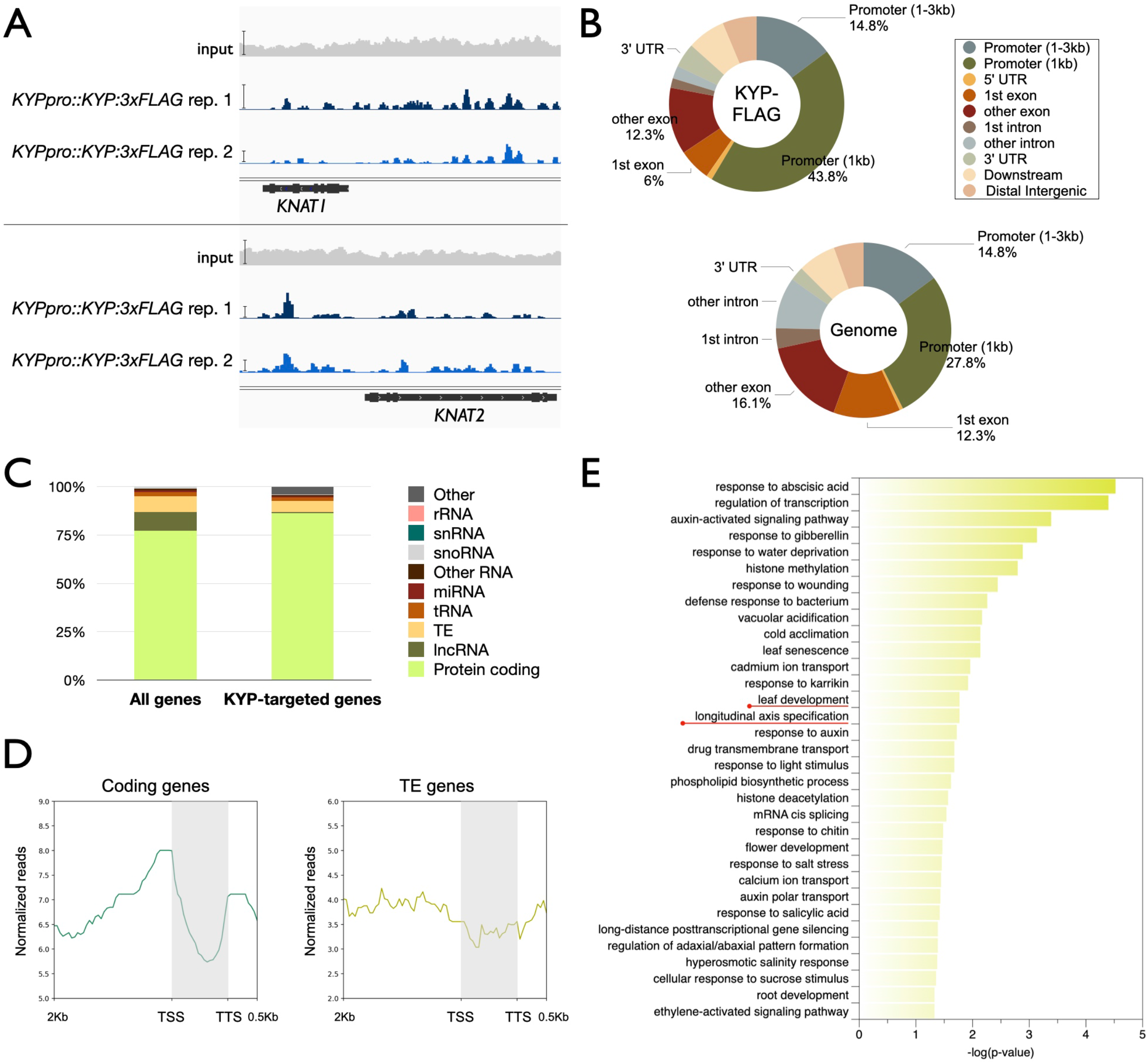
Genome-wide occupancy profile of KYP. (A) Integrated Genome Viewer showing the binding of KYP on *KNAT1* and *KNAT2* genomic regions. Bars: normalized reads = 40. (B) Pie charts showing the distribution of KYP annotated genic and intergenic regions in the genome. (C) Distribution of gene type among all of the KYP-targeted genes. (D) Metagene ChIP-seq binding profiles of KYP among all of the coding genes and TE genes within within the *Arabidopsis* genome. The profile is from 2 kb upstream of the TSS to 0.5 kb downstream of the TTS, and the gene lengths were scaled to the same size. (E) GO-BP annotation of KYP-targeted genes. Annotation terms with *p*-value < 0.05 were listed.

### The genome-wide binding of KYP is positively correlated with AS1 and HDA6

To further analyze the functional correlation between KYP and H3K9me2, we compared the KYP global binding pattern with the H3K9me2 ChIP-seq data of WT and *kyp* from the published dataset (Li at al., 2018). We found that the binding of KYP was more correlated with those genes with their relative H3K9me2 levels are lower in *kyp* compared to WT (Fig. 6A). In contrast, the relative H3 levels in *kyp* compared to WT show no correlation with the binding of KYP (Fig. 6A). Similar results were also obtained when we compared the relative H3K9me2 levels in *kyp/suvh5/6* and WT (Fig. S6). We also found that among those genes showing changed H3K9me2 levels in *kyp*, the general binding pattern of KYP was substantially higher in the genes with decreased H3K9me2 than those genes with increased H3K9me2 level (Fig. 6B). These results indicated that the binding of KYP is indeed correlated with H3K9me2.

**Figure 6.**
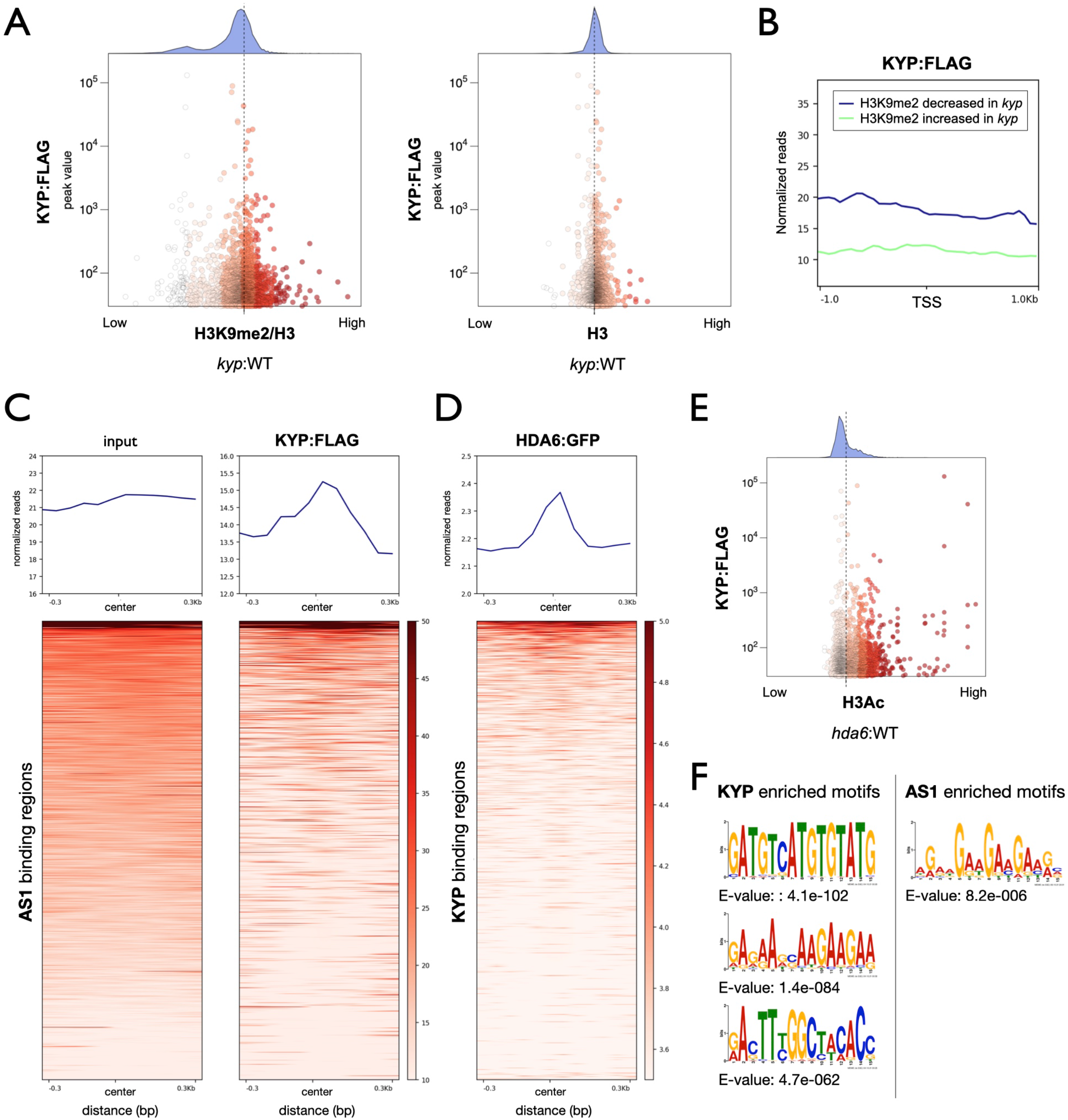
KYP-targeted genomic regions are highly correlated with AS1 and HDA6. (A) X-Y scatter plots showing the relative enrichment of H3K9me2 (*kyp*: WT) and the binding level of KYP. (B) The mean density of KYP enrichment in the gene groups showing decreased- or increased-H3K9me2 in *kyp*. The average KYP enrichment signals within 1 kb genomic regions from up-stream and down-stream of TSS were shown. The KYP enrichment levels were marked in blue (H3K9me2 decreased genes in *kyp*) or green (H3K9me2 increased genes in *kyp*). (C, D) Heat map representation of the co-occupancy of KYP with AS1 (C) and HDA6 (D) in the genome. Each horizontal line represents an AS1-(C) or KYP-(D) bound region, and the signal intensity is shown for KYP (C) or HDA6 (D) binding. Columns show the genomic region surrounding each AS1-(C) or KYP-(D) peak. Signal intensity is indicated by the shade of red. (E) X-Y scatter plots showing the relative enrichment of H3Ac level (*hda6:* WT) and the binding level of KYP. (F) The DNA binding motifs significantly enriched in the KYP or AS1 binding sites.

We also compared the binding of KYP with the previously published chromatin immuno-precipitation with DNA microarray (ChIP-on-chip) data of AS1 (Iwasaki at al., 2013). Plotprofile and plot heatmap analyses indicate that the center of AS1-binding regions was associated with the enrichment of KYP (Fig. 6C), supporting that KYP can be recruited by AS1 to regulate gene expression. In addition, we also compared the KYP-occupied genomic regions with our previously published HDA6 ChIP-seq data (Hung et al., 2020b). We found that the general binding of HDA6 was enriched in the center of KYP-occupied genomic regions (Fig. 6D), supporting that HDA6 interacts with KYP to synergistically regulate gene expression. Furthermore, we also compared the KYP global binding pattern with the H3Ac ChIP-seq data of WT and *hda6* from the published dataset (Hung et al., 2020b). The binding of KYP was more correlated with the genes with their relative H3Ac levels are increased in *hda6* compared to WT (Fig. 6E). Collectively, these results suggested that the binding of KYP is associated with the H3Ac changes regulated by HDA6.

We further identified the cis-elements including “GATGTCATGTGTATG”, “RACTTYGGCTACACC” and (AG/AAG)_n_ repeat sequences that were enriched within the KYP binding sites, (Fig. 6F). Interestingly, the (AG/AAG)_n_ repeat was also found in the AS1-occupied genomic regions identified previously by ChIP-on-chip (Fig. 6F). In addition, previous published data also shown that the (AG/AAG)_n_ repeat was enriched in the HDA6 occupied genomic regions (Hung et al., 2020b). Together, these results support that KYP can cotarget on the similar genomic regions with HDA6 and AS1.

### KYP and HDA6 co-target a subset of leaf development genes

In addition to *KNAT1* and *KNAT2*, we also identified additional genes that were targeted by KYP, such as *KNAT3*, *KNAT5*, *NUCLEOLIN 1* (*NUC1*), *GROWTH-REGULATING FACTOR 4* (*GRF4*) and *CYCLIN DEPENDENT PROTEIN KINASE 2* (*CDKC2*) (Fig. 7A). The class II *KNOX* genes *KNAT3* and *KNAT5* are involved in development of all above-ground organs in *Arabidopsis* and *knat3/4/5* mutant plants display developmental defected leaves (Furumizu et al., 2015). *Arabidopsis* NUC1 is a nucleolin protein that is involved in rRNA processing, ribosome biosynthesis, and vascular pattern formation (Matsumura et al. 2016). NUC1 is also involved in leaf development and is functional associated with AS2 (Vial-Pradel et al. 2018). The transcription factor GRF4 and the cell cycle regulator CDKC2 were also found to be involved in the regulation of leaf development (Kuijt et al., 2014; Zhao et al., 2017). In addition to leaf development, these KYP target genes including *NUC1, GRF4* and *MLP328* are also involved in other biological processes, such as flowering, root development, stress responses and cell well formation (Kuang et al., 2009; Guo et al., 2011; Wang et al., 2015; Prado et al., 2019; Jiang et al., 2020; Wang et al., 2020). Genome browser views of the KYP ChIP-Seq data indicate that the KYP enriched regions on these genes were highly correlated with the HDA6 enriched regions (Fig. 7A). The binding of KYP on these genes was further confirmed by ChIP-qPCR (Fig. 7B). qRT-PCR analyses indicate that the expression of these KYP-HDA6 co-targeted genes was significantly increased in the *hda6/kyp/suvh5/6* quadruple mutant compared to WT (Fig. 7C). Collectively, these results indicate that KYP co-targets a subset of genes with HDA6 to regulate leaf development.

**Figure 7.**
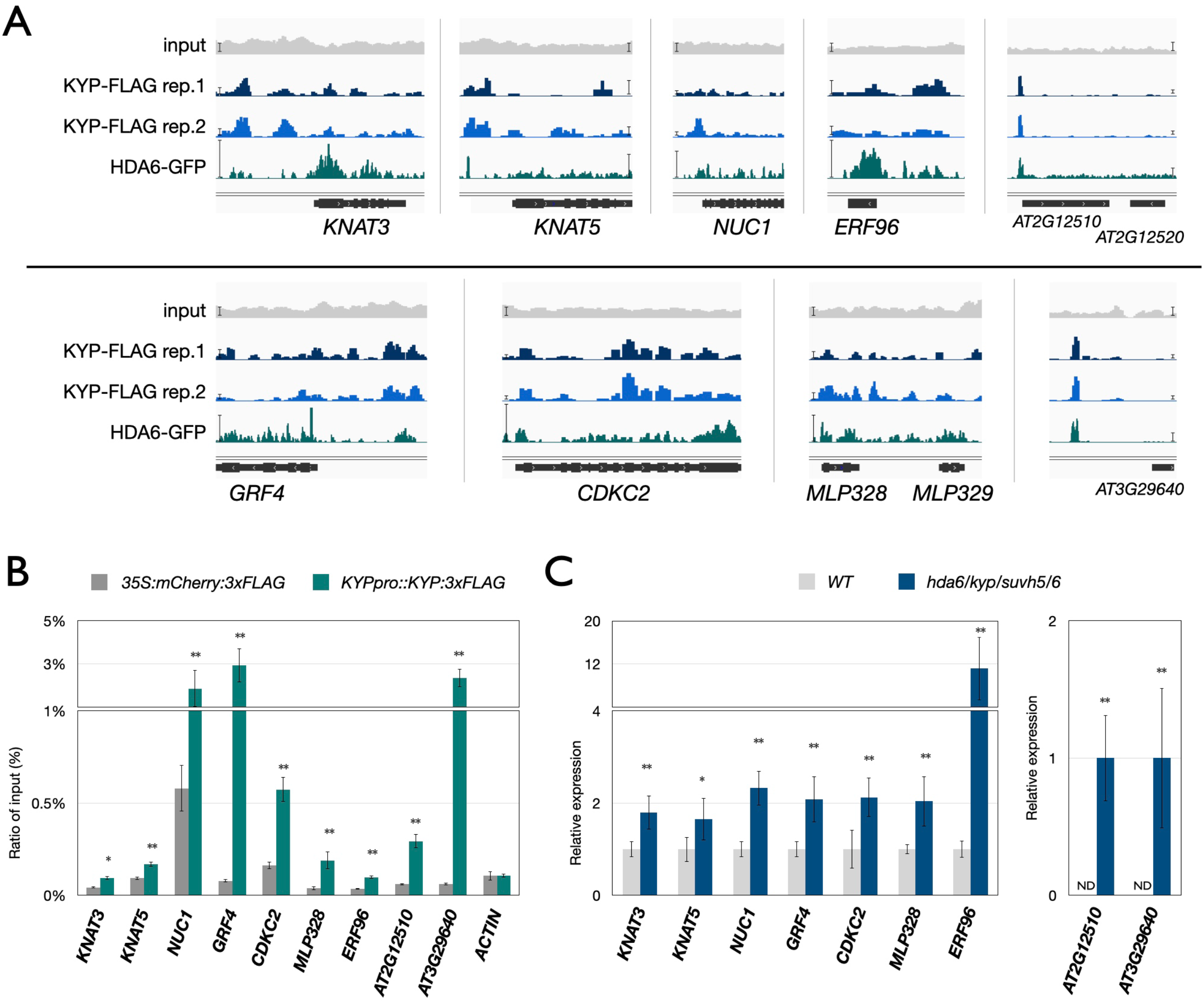
The binding and expression of KYP regulated genes. (A) Integrated Genome Viewer showing the binding of KYP and HDA6 in the selected genes. Bars: normalized reads = 10. (B, C) Binding of KYP (B) and gene expression patterns (C) in the indicated samples. ChIP assays were performed with the anti-FLAG antibody. The amount of immunoprecipitated DNA was quantified by qPCR. Values represent the normalized average immunoprecipitation effi-ciencies (%) against the total input DNA. *UBQ10* was used as an internal control. ND: not detected. Error bars correspond to SD. *: P< 0.05, **: P< 0.005 (Student’s t-test). At least three independent biological replicates were performed with similar results.

## Discussion

In *Arabidopsis*, 31 SDG proteins predicted to have HKMT activity can be further classified into five classes (class I to class V) based on their domain architectures or their target lysine residues (Ng et al., 2007). There are 15 Class V SDG proteins including 10 SUVH proteins and 5 SUVR proteins in *Arabidopsis*. Several Class V SDG proteins have been found to be associated with H3K9 methylation involved in heterochromatin maintenance and DNA methylation (Pontes et al., 2006; Grafi et al., 2007; Johnson et al., 2007; Pontvianne et al., 2010). SUVHs contain an N-terminal SRA domain and a SET domain in C-terminal (Baumbusch et al., 2001; Johnson et al., 2007; Rajakumara et al., 2011). The SRA domain is required for direct binding to methylated DNA (Johnson et al., 2007; Rajakumara et al., 2011). KYP, SUVH5, and SUVH6, the best characterized SUVH proteins in *Arabidopsis*, are H3K9me1/2 methyltransferases responsible for chromatin silencing (Jackson et al., 2002; Ebbs and Bender, 2006; Stroud et al., 2014). Two other SUVH proteins, SUVH2 and SUVH9, are inactive for histone methyltransferase activity, but they can recruit RNA polymerase V to chromatin by associating with the DDR (DRD1 peptide-DMS3-RDM1) complex (Johnson et al., 2014; Liu et al., 2014). In addition, the SUVR proteins SUVR4 and SUVR5 have been found to be involved in H3K9me *in vivo* (Thorstensen et al., 2006; Veiseth et al., 2011; Caro et al., 2012). Collectively, these studies indicate that the Class V SDG proteins are important in gene silencing by regulating H3K9me.

H3K9me2 is a crucial histone modification marker during embryo-development in both plant and mammalian systems (Graf et al., 2014; Au Yeung et al., 2019; Parent et al., 2021). Recent studies have also showed that H3K9me2 is important in regulating gene expression involved in *Arabidopsis* development (Saze et al., 2008; Dutta et al., 2017; Hung et al., 2020a; Xu and Jiang, 2020; Hung et al., 2021). Although KYP and SUVH5/6 have been identified as crucial regulators of H3K9me2 in *Arabidopsis*, the function of KYP and SUVH5/6 in plant development remains elusive. In the present study, we found that KYP and SUVH5/6 are functionally associated with HDA6. Furthermore, HDA6 and KYP/SUVH5/6 function synergistically to regulate the core leaf development genes including *KNAT1* and *KNAT2*. A recent study also demonstrated that another Class V SDG protein SUVH9 is involved in embryonic development by regulating asymmetric DNA methylation (Parent et al., 2021). Taken together, these results indicate that the Class V SDG proteins including KYP and SUVH5/6 play important roles in plant developmental processes.

*KNAT1* and *KNAT2* are Class I *KNOX* homeobox genes and play important roles in meristem development and leaf morphogenesis (Chuck et al., 1996; Venglat et al., 2002; Byrne et al., 2005). Previous studies have demonstrated that the expression of *KNAT1* and *KNAT2* is associated with the changes in H3Ac, H3K9me2 and H3K27me3 (Luo et al., 2012; Lodha et al., 2013; You et al., 2017). In this study, we found that KYP/SUVH5/6 and HDA6 function synergistically to regulate *KNAT1* and *KNAT2* by altering H3K9me2 and H3Ac levels. Furthermore, the expression of the *KNOX* genes was increased in the *hda6/kyp/suvh5/6* quadruple mutant compared to *hda6* and *kyp/suvh5/6*. Similarly, the expression of TEs was also increased in the *hda6/kyp/suvh5/6* quadruple mutant compared to *hda6* or *kyp/suvh5/6* (Yu et al., 2017). These results suggest that both of the H3K9me2 decrease and H3Ac increase are required to activate expression of the protein coding genes. Interestingly, it has been shown that there is an antagonistic pattern of H3K9me2 and H3Ac enrichment during embryogenesis in both plant and mammalian systems (Lee et al., 2012; Rodriguez-Sanz et al., 2014), indicating a functional crosstalk between H3K9me2 and H3Ac in developmental processes. Recent studies demonstrated that *Arabidopsis* HDA6 is also functionally associated with the H3K4 demethylases LDL1/2 and FLD (Yu et al., 2011; Hung et al., 2018; Hung et al., 2019; Hung et al., 2020b). It remains to be determined whether KYP/SUVH5/6 are also functionally associated with H3K4 demethylases.

In this study, we found that KYP and SUVH5/6 can direct interact with AS1-AS2 to regulate the expression of *KNAT1/2* by altering H3Ac and H3K9me2 levels. Accumulation of H3K9me2 is highly associated with DNA methylation on CHG and CHH sites (Grafi et al., 2007; Johnson et al., 2007; Pontes et al., 2006; Pontvianne et al, 2010). Interestingly, AS1-AS2 is also required for maintaining DNA methylation on *ETTIN/AUXIN RESPONSE FAC-TOR3* (*ETT/ARF3*) (Iwasaki at al., 2013; Vial-Pradel et al. 2018), suggesting that AS1-AS2 and KYP/SUVH5/6 may also function together in the regulation of DNA methylation. Recent studies indicated that AS2 is highly associated with chromocenter including ribosomal DNA repeat regions and is involved in the regulation of cell division (Luo et al., 2020; Machida et al., 2021). It remains to be determined whether KYP/SUVH5/6 are also involved in these processes.

In addition to *KNAT1/2*, other leaf development genes including *KNAT3, KNAT5, NUC1, GRF4* and *CDKC2* (Furumizu et al., 2015; Matsumura et al. 2016; Vial-Pradel et al. 2018; Kuijt et al., 2014; Zhao et al., 2017) are also regulated by KYP/SUVH5/6-HDA6. The GO-BP analysis indicates that KYP targeted genes are associated with stress responses, hormone responses and different developmental processes. It has been shown that KYP is involved in regulating seed dormancy by repressing ABA signaling genes (Zheng et al., 2012). Furthermore, SUVH5 has been found to act as a positive regulator of light-mediated seed germination (Gu et al., 2019). Interestingly, we found that the GO-terms “response to abscisic acid” and “response to light stimulus” were also enriched in KYP targeted genes. Taken together, these results indicate that KYP/SUH5/6 are involved in various developmental processes and pathways. In the genome-wide occupancy profile of KYP, we found that the binding of KYP was more enriched in promoter regions, and most of the KYP targeted genes are protein coding genes. Furthermore, the binding of KYP is highly correlated with the binding of AS1 and HDA6. Together, these data support the notion that AS1/2 recruit the transcriptional repression complex containing HDA6 and KYP/SUVH5/6 to regulate gene expression.

In conclusion, this study provides insight about how the H3K9 methyltransferases KYP and SUVH5/6 are involved in leaf development by interacting with AS1-AS2 (Fig. 8). The AS1-AS2 transcription complex acts as transcription repressor by recruiting KYP/SUVH5/6-HDA6 histone modification complex to repress the expression the *KNOX* genes *KNAT1* and *KNAT2* via H3K9me2 and H3 deacetylation. In addition, the KYP/SUVH5/6-HDA6 histone modification complex can also regulate gene expression involved in other developmental processes and pathways.

**Figure 8.**
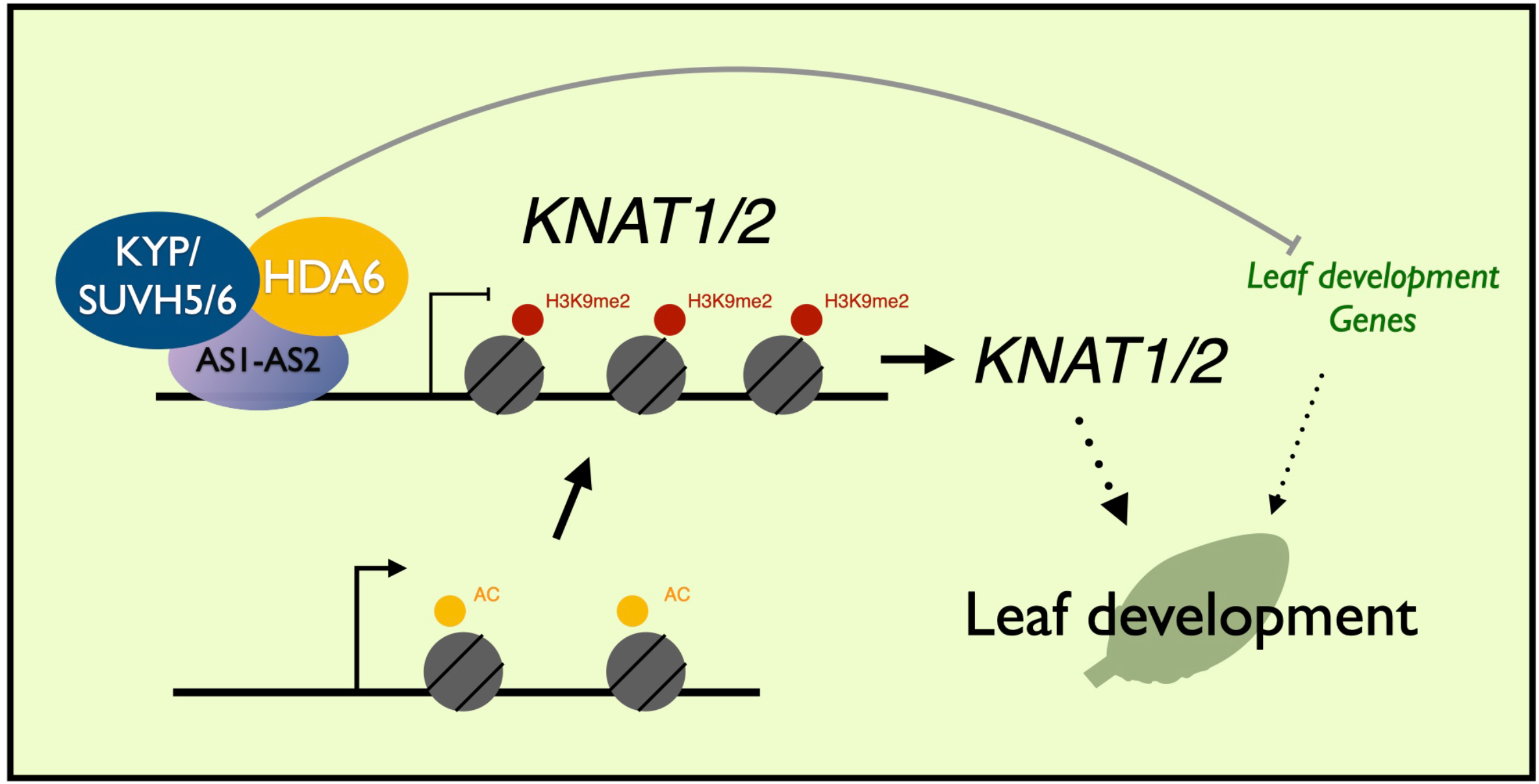
A model for the KYP/SUVH5/6 function in the regulation of the *KNAT1/2* and leaf development genes. *Arabidopsis* KYP/SUVH5/6 are involved in leaf development by repressing *KNAT1/2*. AS1-AS2 act as transcription repressors and recruit HDA6-KYP/SUVH5/6 to *KNAT1/2by altering* H3Ac/H3K9me2 levels. KYP/SUVH5/6 play an important role in leaf development by regulating the expression of *KNAT1/2* and other leaf development genes.

## Materials and Methods

### Plant materials and growth conditions

*Arabidopsis* (*Arabidopsis thaliana*) plants were germinated and grown in 22°C under long day (LD) (16 h light /8 h dark cycle) conditions. *kyp/suvh4-3*, (SALK_130630), *suvh5* (GABI_263C05), *suvh6* (SAIL_1244_F04), *hda6-6* (*axe1-5*) as well as the *kyp/suvh5/suvh6* (*kyp/suvh5/6*) triple mutant and the *hda6/kyp/suvh5/suvh6* (*hda6/kyp/suvh5/6*) quadruple mutant were reported previously (Murfett et al., 2001; Johnson et al., 2008; Zheng et al., 2012; Yu et al., 2017). The *hda6/kyp* double mutant was generated by crossing *kyp* (*suvh4-3*) and *hda6-6* (*axe1-5*). All mutants used in this study are in the Col-0 background.

### Plasmid construction and plant transformation

The full-length coding sequences (CDS) of *KYP*, *SUVH5*, *SUVH6*, *AS1* and *AS2* were used in the previous published studies (Luo et al., 2012; Yu et al., 2017). To generate the *KYPpro::KYP:GFP* and *KYPpro::KYP:3xFLAG* constructs, the 5 kb *KYP* genomic DNA sequence containing the 2 kb *KYP* native promoter was PCR-amplified and cloned into the *pCR8/GW/TOPO* vector (Invitrogen), then recombined into a modified *pEarleyGate302* vector containing the *3xFLAG* tag or *PMDC107* vector with the *mGFP* tag.

*KYPpro::KYP:GFP/kyp* and *KYPpro::KYP:3xFLAG/kyp* transgenic plants were generated by transforming *KYPpro::KYP:GFP* or *KYPpro::KYP:3xFLAG* into the *kyp* mutant by the floral dip method. To express *KYPpro::KYP:3xFLAG* in the *as1* mutant background, *KYP-pro::KYP:3xFLAG* plants were crossed with the *as1* mutant.

### Bimolecular fluorescence complementation and co-immunoprecipitation assays

Bimolecular fluorescence complementation (BiFC) assays were performed as previously described (Hung et al., 2018). To generate the constructs for BiFC assays, full-length or truncated cDNA fragments of *KYP*, *SUVH5*, *SUVH6*, *AS1* and *AS2* were PCR-amplified and cloned into the *pCR8/GW/TOPO* vector (Invitrogen), and then recombined into the YN vector *pEarleyGate201-YN* and the YC vector *pEarleyGate202-YC*. Constructed vectors were transiently transformed into *Arabidopsis* protoplasts or tobacco (*Nicotiana benthamiana*) leaves. Transfected protoplasts or leaves were then examined by using a TCS SP5 confocal spectral microscope imaging system (Leica, https://www.leica.com/).

Co-immunoprecipitation assays were performed as previously described (Hung et al., 2018). Anti-GFP (Santa Cruz Biotechnologies, catalog no. SC-9996; 1:3000 dilution) and anti-AS1 (SIGMA catalog no. M2; 1:3000 dilution) antibodies were used as primary antibodies for Western blot, the resulting signals were detected by using a Pierce ECL Western blotting kit (Pierce, https://www.lifetechnologies.com/).

### Quantitative reverse transcriptase PCR analysis

Total RNA was isolated using TRIZOL reagent (Invitrogen, 15596026) according to the manufacturer’s instructions. Two micrograms of total RNA treated with DNAse (Promega, RQ1 #M6101) were used to synthesize cDNA (Promega, #1012891). RT-qPCR (Real-Time quantitative PCR) was performed using iQ SYBR Green Supermix solution (Bio-Rad, #170-8880). The CFX96 Real-Time PCR Detection System (Bio-Rad Laboratories, Inc.) was used with the following cycling conditions: 95°C for 10 min, followed by 45 cycles of 95°C for 15 s, 60°C for 30s, and then fluorescent detection. This was immediately followed by a melting curve analysis (65–95°C, incrementing 0.5°C for 5 s, and plate reading) to confirm the absence of non-specific products. Each sample was quantified at least in triplicate, and normalized by calculating delta Cq (quantification cycle) to the expression of the internal control *Ubiquitin10* (*UBQ10*). The Cq and relative expression level are calculated by the Biorad CFX Manager 3.1 based on the MIQE guidelines. Standard deviations represent at least 3 technical and 2 biological replicates. The variance in average data is represented by SEM (standard error of the mean). The SD (standard deviation), SEM determination and P-value were calculated using Student’s paired t-test. The gene specific primers used for qRT-PCR are listed in Table S1.

### Chromatin immunoprecipitation assays

ChIP assays were performed as previously described (Hung et al., 2018). Chromatin extracts were prepared from seedlings treated with 1% formaldehyde. Chromatin was sheared to the mean length of 500 bp by sonication, proteins and DNA fragments were then immuno-precipitated using antibodies against anti-FLAG (SIGMA, catalog no. M2), H3Ac (Millipore, catalog no. 06-599) or H3K9me2 (diagenode, C15410060). The DNA cross-linked to immunoprecipitated proteins were reversed, and then analyzed by real-time PCR using specific primers (Table S1). Percent input was calculated as follows: 2∧(Cq(IN)-Cq(IP))X100. Cq is the quantification cycle as calculated by the Biorad CFX Manager 3.1 based on the MIQE guidelines. Standard deviations represent at least 3 technical and 2 biological replicates. The variance in average data is represented by SEM (standard error of the mean). The SD (standard deviation), SEM determination and P-value were calculated using Student’s paired t-test.

### ChIP-seq and data analyses

ChIP-seq assays were performed based on previous research (Hung et al., 2018, Hung et al., 2020b). 2 ng of DNA from ChIP was pooled to ensure that there are enough starting DNA for library construction. Two biological replicates were prepared and sequenced for each ChIP-seq experiment. The ChIP DNA was first tested by qRT-PCR and then used to prepare ChIP-seq libraries. End repair, adaptor ligation, and amplification were carried out using the NEBNext^®^ Ultra^™^ II DNA Library Prep kit (cat no. E7645) according to the manufacturer’s protocol. The Novoseq PE150 was used for high-throughput sequencing of the ChIP-seq libraries. The raw sequence data were processed using the GAPipeline Illumina sequence data analysis pipeline. Bowtie2 was then employed to map the reads to the *Arabidopsis* genome (TAIR10) (Lamesch et al., 2012). Only perfectly and uniquely mapped reads were retained for further analysis. The alignments were first converted to Wiggle (WIG) files using deepTools. The data were then imported into the Integrated Genome Viewer (IGV) (Thorvaldsdottir et al., 2013) for visualization. The distribution of the ChIP binding peaks was analyzed with ChIPseeker (Table S2) (Yu et al., 2015), and a high-read random *Arabidopsis* genomic region subset (1,350,000 regions) was used to represent the ratio of the total *Arabidopsis* genomic regions. To identify DNA motifs enriched sites, 400-bp sequences encompassing each peak summit (200 bp upstream and 200 bp downstream) were extracted and searched for enriched DNA motifs using MEME-ChIP with the default parameters (Machanick and Bailey, 2011).

The KYP:FLAG ChIP-seq short read data have been submitted to the NCBI Gene Expression Omnibus (GEO) database (GSE195735).

## Acknowledgements

We thank Technology Commons, College of Life Science, National Taiwan University for the convenient use of the Apatome 2.0 microscope, TCS SP5 confocal spectral microscope imaging system and the Bio-Rad real-time PCR system.

## Funding

This work is supported by the Ministry of Science and Technology of the Republic of China (108-2311-B-002-013-MY3 and 110-2311-B-002-027 to K. W.) and National Taiwan University (111L893001 to K.W.).

## Authors’ Contributions

F.-Y. H., and K. W. designed research, Y.-R. F., F.-Y. H., Y.-C. L. and Y.-H. S. performed research. F.-Y. H., Y.-R. F., and K. W. analyzed data. F.-Y. H., Y.-R. F., and K. W. wrote the article.

## Parsed Citations

Au Yeung WK, Brind’Amour J, Hatano Y, Yamagata K, Feil R, Lorincz MC, Tachibana M, Shinkai Y, Sasaki H. 2019. Histone H3K9 methyltransferase G9a in oocytes Is essential for preimplantation development but dispensable for CG methylation protection. Cell Rep 27: 282–293 e284. Google Scholar: <u>Author Only Title Only Author and Title</u>

Baumbusch LO, Thorstensen T, Krauss V, Fischer A, Naumann K, Assalkhou R, Schulz I, Reuter G, Aalen RB. 2001. The Arabidopsis thaliana genome contains at least 29 active genes encoding SET domain proteins that can be assigned to four evolutionarily conserved classes. Nucleic Acids Res 29: 4319–4333. Google Scholar: <u>Author Only Title Only Author and Title</u>

Berger SL. 2007. The complex language of chromatin regulation during transcription. Nature 447: 407–412. Google Scholar: <u>Author Only Title Only Author and Title</u>

Bernatavichute YV, Zhang X, Cokus S, Pellegrini M, Jacobsen SE. 2008. Genome-wide association of histone H3 lysine nine methylation with CHG DNA methylation in Arabidopsis thaliana. PLoS One 3: e3156. Google Scholar: <u>Author Only Title Only Author and Title</u>

Byrne ME. 2005. Networks in leaf development. Current Opinion in Plant Biology 8: 59–66. Google Scholar: <u>Author Only Title Only Author and Title</u>

Byrne ME, Barley R, Curtis M, Arroyo JM, Dunham M, Hudson A, Martienssen RA. 2000. Asymmetric leaves1 mediates leaf patterning and stem cell function in Arabidopsis. Nature 408: 967–971. Google Scholar: <u>Author Only Title Only Author and Title</u>

Caro E, Stroud H, Greenberg MV, Bernatavichute YV, Feng S, Groth M, Vashisht AA, Wohlschlegel J, Jacobsen SE. 2012. The SET-domain protein SUVR5 mediates H3K9me2 deposition and silencing at stimulus response genes in a DNA methylation-independent manner. PLoS Genet 8: e1002995. Google Scholar: <u>Author Only Title Only Author and Title</u>

Chuck G, Lincoln C, Hake S. 1996. KNAT1 induces lobed leaves with ectopic meristems when overexpressed in Arabidopsis. Plant Cell 8: 1277–1289. Google Scholar: <u>Author Only Title Only Author and Title</u>

Du J, Johnson LM, Groth M, Feng S, Hale CJ, Li S, Vashisht AA, Gallego-Bartolome J, Wohlschlegel JA, Patel DJ et al. 2014. Mechanism of DNA methylation-directed histone methylation by KRYPTONITE. Molecular Cell 55: 495–504. Google Scholar: <u>Author Only Title Only Author and Title</u>

Du J, Johnson LM, Jacobsen SE, Patel DJ. 2015. DNA methylation pathways and their crosstalk with histone methylation. Nature Reviews Molecular Cell Biology 16: 519–532. Google Scholar: <u>Author Only Title Only Author and Title</u>

Du J, Zhong X, Bernatavichute YV, Stroud H, Feng S, Caro E, Vashisht AA, Terragni J, Chin HG, Tu A et al. 2012. Dual binding of chromomethylase domains to H3K9me2-containing nucleosomes directs DNA methylation in plants. Cell 151: 167–180. Google Scholar: <u>Author Only Title Only Author and Title</u>

Dutta A, Choudhary P, Caruana J, Raina R. 2017. JMJ27, an Arabidopsis H3K9 histone demethylase, modulates defense against Pseudomonas syringae and flowering time. Plant Journal 91: 1015–1028. Google Scholar: <u>Author Only Title Only Author and Title</u>

Ebbs ML, Bender J. 2006. Locus-specific control of DNA methylation by the Arabidopsis SUVH5 histone methyltransferase. Plant Cell 18: 1166–1176. Google Scholar: <u>Author Only Title Only Author and Title</u>

Furumizu C, Alvarez JP, Sakakibara K, Bowman JL. 2015. Antagonistic roles for KNOX1 and KNOX2 genes in patterning the land plant body plan following an ancient gene duplication. Plos Genetics 11: e1004980. Google Scholar: <u>Author Only Title Only Author and Title</u>

Grafi G, Ben-Meir H, Avivi Y, Moshe M, Dahan Y, Zemach A. 2007. Histone methylation controls telomerase-independent telomere lengthening in cells undergoing dedifferentiation. Developmental Biology 306: 838–846. Google Scholar: <u>Author Only Title Only Author and Title</u>

Gu D, Ji R, He C, Peng T, Zhang M, Duan J, Xiong C, Liu X. 2019. Arabidopsis histone methyltransferase SUVH5 Is a positive regulator of light-mediated seed germination. Front Plant Sci 10: 841. Google Scholar: <u>Author Only Title Only Author and Title</u>

Guo D, Wong WS, Xu WZ, Sun FF, Qing DJ, Li N. 2011. Cis-cinnamic acid-enhanced 1 gene plays a role in regulation of Arabidopsis bolting. Plant Mol Biol 75: 481–495. Google Scholar: <u>Author Only Title Only Author and Title</u>

Guo M, Thomas J, Collins G, Timmermans MCP. 2008. Direct repression of KNOX loci by the ASYMMETRIC LEAVES1 complex of Arabidopsis. Plant Cell 20: 48–58. Google Scholar: <u>Author Only Title Only Author and Title</u>

Hung FY, Chen FF, Li C, Chen C, Lai YC, Chen JH, Cui Y, Wu K. 2018. The Arabidopsis LDL1/2-HDA6 histone modification complex is functionally associated with CCA1/LHY in regulation of circadian clock genes. Nucleic Acids Res 46: 10669–10681. Google Scholar: <u>Author Only Title Only Author and Title</u>

Hung FY, Chen FF, Li C, Chen C, Chen JH, Cui Y, Wu K. 2019. The LDL1/2-HDA6 histone modification complex interacts with TOC1 and regulates the core circadian clock components in Arabidopsis. Front Plant Sci 10: 233. Google Scholar: <u>Author Only Title Only Author and Title</u>

Hung F-Y, Chen J-H, Feng Y-R, Lai Y-C, Yang S, Wu K. 2020a. Arabidopsis JMJ29 is involved in trichome development by regulating the core trichome initiation gene GLABRA3. Plant Journal 103: 1735–1743. Google Scholar: <u>Author Only Title Only Author and Title</u>

Hung F-Y, Chen C, Yen M-R, Hsieh J-WA, Li C, Shih Y-H, Chen F-F, Chen P-Y, Cui Y, Wu K. 2020b. The expression of long noncoding RNAs is associated with H3Ac and H3K4me2 changes regulated by the HDA6-LDL1/2 histone modification complex in Arabidopsis. Nar Genomics and Bioinformatics 2: lqaa066. Google Scholar: <u>Author Only Title Only Author and Title</u>

Hung F-Y, Lai Y-C, Wang J, Feng Y-R, Shih Y-H, Chen J-H, Sun H-C, Yang S, Li C, Wu K. 2021. The Arabidopsis histone demethylase JMJ28 regulates CONSTANS by interacting with FBH transcription factors. Plant Cell 33: 1196–1211. Google Scholar: <u>Author Only Title Only Author and Title</u>

Iwakawa H, Ueno Y, Semiarti E, Onouchi H, Kojima S, Tsukaya H, Hasebe M, Soma T, Ikezaki M, Machida C et al. 2002. The ASYMMETRIC LEAVES2 gene of Arabidopsis thaliana, required for formation of a symmetric flat leaf lamina, encodes a member of a novel family of proteins characterized by cysteine repeats and a leucine zipper. Plant Cell Physiol 43: 467–478. Google Scholar: <u>Author Only Title Only Author and Title</u>

Iwasaki M, Takahashi H, Iwakawa H, Nakagawa A, Ishikawa T, Tanaka H, Matsumura Y, Pekker I, Eshed Y, Vial-Pradel S et al. 2013. Dual regulation of ETTIN (ARF3) gene expression by AS1-AS2, which maintains the DNA methylation level, is involved in stabilization of leaf adaxial-abaxial partitioning in Arabidopsis. Development 140: 1958–1969. Google Scholar: <u>Author Only Title Only Author and Title</u>

Jackson D, Veit B, Hake S. 1994. Expression of maize KNOTTED1 related homeobox genes in the shoot apical meristem predicts patterns of morphogenesis in the vegetative shoot. Development 120: 405–413. Google Scholar: <u>Author Only Title Only Author and Title</u>

Jackson JP, Lindroth AM, Cao XF, Jacobsen SE. 2002. Control of CpNpG DNA methylation by the KRYPTONITE histone H3 methyltransferase. Nature 416: 556–560. Google Scholar: <u>Author Only Title Only Author and Title</u>

Jiang L, Yang J, Liu C, Chen Z, Yao Z, Cao S. 2020. Overexpression of ethylene response factor ERF96 gene enhances selenium tolerance in Arabidopsis. Plant Physiol Biochem 149: 294–300. Google Scholar: <u>Author Only Title Only Author and Title</u>

Johnson LM, Bostick M, Zhang X, Kraft E, Henderson I, Callis J, Jacobsen SE. 2007. The SRA methyl-cytosine-binding domain links DNA and histone methylation. Current Biology 17: 379–384. Google Scholar: <u>Author Only Title Only Author and Title</u>

Kidner CA, Timmermans MCP, Byrne ME, Martienssen RA 2002. Developmental genetics of the angiosperm leaf. Adv Bot Res 38: 191–234. Google Scholar: <u>Author Only Title Only Author and Title</u>

Klose RJ, Yi Z. 2007. Regulation of histone methylation by demethylimination and demethylation. Nature Reviews Molecular Cell Biology 8: 307–318. Google Scholar: <u>Author Only Title Only Author and Title</u>

Kuijt SJ, Greco R, Agalou A. Shao J, t Hoen CC, Overnas E, Osnato M, Curiale S, Meynard D, van Gulik R et al. 2014. Interaction between the GROWTH-REGULATING FACTOR and KNOTTED1-LIKE HOMEOBOX families of transcription factors. Plant Physiol 164: 1952–1966. Google Scholar: <u>Author Only Title Only Author and Title</u>

Lamesch P, Berardini TZ, Li D, Swarbreck D, Wilks C, Sasidharan R, Muller R, Dreher K, Alexander DL, Garcia-Hernandez M et al. 2012. The Arabidopsis Information Resource (TAIR): improved gene annotation and new tools. Nucleic Acids Res 40: D1202–1210. Google Scholar: <u>Author Only Title Only Author and Title</u>

Lee HA, Cho HM, Lee DY, Kim KC, Han HS, Kim IK. 2012. Tissue-specific upregulation of angiotensin-converting enzyme 1 in spontaneously hypertensive rats through histone code modifications. Hypertension 59: 621–626. Google Scholar: <u>Author Only Title Only Author and Title</u>

Li X, Harris CJ, Zhong Z, Chen W, Liu R, Jia B, Wang Z, Li S, Jacobsen SE, Du J. 2018. Mechanistic insights into plant SUVH family H3K9 methyltransferases and their binding to context-biased non-CG DNA methylation. Proceedings of the National Academy of Sciences of the United States of America 115: E8793–E8802. Google Scholar: <u>Author Only Title Only Author and Title</u>

Liu C, Lu F, Cui X, Cao X. 2010. Histone methylation in higher plants. in Annual Review of Plant Biology, pp. 395–420. Google Scholar: <u>Author Only Title Only Author and Title</u>

Liu X, Yu C-W, Duan J, Luo M, Wang K, Tian G, Cui Y, Wu K. 2012. HDA6 directly interacts with DNA methyltransferase MET1 and maintains transposable element silencing in Arabidopsis. Plant Physiology 158: 119–129. Google Scholar: <u>Author Only Title Only Author and Title</u>

Lodha M, Marco CF, Timmermans MC. 2013. The ASYMMETRIC LEAVES complex maintains repression of KNOX homeobox genes via direct recruitment of Polycomb-repressive complex2. Genes Dev 27: 596–601. Google Scholar: <u>Author Only Title Only Author and Title</u>

Long JA, Moan EI, Medford JI, Barton MK. 1996. A member of the KNOTTED class of homeodomain proteins encoded by the STM gene of Arabidopsis. Nature 379: 66–69. Google Scholar: <u>Author Only Title Only Author and Title</u>

Luo L, Ando S, Sakamoto Y, Suzuki T, Takahashi H, Ishibashi N, Kojima S, Kurihara D, Higashiyama T, Yamamoto KT et al. 2020. The formation of perinucleolar bodies is important for normal leaf development and requires the zinc-finger DNA-binding motif in Arabidopsis ASYMMETRIC LEAVES2. Plant J 101: 1118–1134. Google Scholar: <u>Author Only Title Only Author and Title</u>

Luo M, Yu C-W, Chen F-F, Zhao L, Tian G, Liu X, Cui Y, Yang J-Y, Wu K. 2012. Histone deacetylase HDA6 is functionally associated with AS1 in repression of KNOX genes in Arabidopsis. Plos Genetics 8: e1003114. Google Scholar: <u>Author Only Title Only Author and Title</u>

Machanick P, Bailey TL. 2011. MEME-ChIP: motif analysis of large DNA datasets. Bioinformatics 27: 1696–1697. Google Scholar: <u>Author Only Title Only Author and Title</u>

Machida Y, Suzuki T, Sasabe M, Iwakawa H, Kojima S, Machida C. 2021. Arabidopsis ASYMMETRIC LEAVES2 (AS2): roles in plant morphogenesis, cell division, and pathogenesis. J Plant Res. Google Scholar: <u>Author Only Title Only Author and Title</u>

Magnani E, Hake S. 2008. KNOX lost the OX: The Arabidopsis KNATM gene defines a novel class of KNOX transcriptional regulators missing the homeodomain. Plant Cell 20: 875–887. Google Scholar: <u>Author Only Title Only Author and Title</u>

Matsumura Y, Ohbayashi I, Takahashi H, Kojima S, Ishibashi N, Keta S, Nakagawa A, Hayashi R, Saez-Vasquez J, Echeverria M et al. 2016. A genetic link between epigenetic repressor AS1-AS2 and a putative small subunit processome in leaf polarity establishment of Arabidopsis. Biol Open 5: 942–954. Google Scholar: <u>Author Only Title Only Author and Title</u>

Murfett J, Wang XJ, Hagen G, Guilfoyle TJ. 2001. Identification of Arabidopsis histone deacetylase HDA6 mutants that affect transgene expression. Plant Cell 13: 1047–1061. Google Scholar: <u>Author Only Title Only Author and Title</u>

Ng DWK, Wang T, Chandrasekharan MB, Aramayo R, Kertbundit S, Hall TC. 2007. Plant SET domain-containing proteins: Structure, function and regulation. Biochimica Et Biophysica Acta-Gene Structure and Expression 1769: 316–329. Google Scholar: <u>Author Only Title Only Author and Title</u>

Ori N, Eshed Y, Chuck G, Bowman JL, Hake S. 2000. Mechanisms that control knox gene expression in the Arabidopsis shoot. Development 127: 5523–5532. Google Scholar: <u>Author Only Title Only Author and Title</u>

Parent JS, Cahn J, Herridge RP, Grimanelli D, Martienssen RA. 2021. Small RNAs guide histone methylation in Arabidopsis embryos. Genes Dev 35: 841–846. Google Scholar: <u>Author Only Title Only Author and Title</u>

Pontes O, Li CF, Nunes PC, Haag J, Ream T, Vitins A, Jacobsen SE, Pikaard CS. 2006. The Arabidopsis chromatin-modifying nuclear siRNA pathway involves a nucleolar RNA processing center. Cell 126: 79–92. Google Scholar: <u>Author Only Title Only Author and Title</u>

Pontvianne F, Blevins T, Pikaard CS. 2010. Arabidopsis histone lysine methyltransferases. in Adv Bot Res, pp. 1–22. Google Scholar: <u>Author Only Title Only Author and Title</u>

Prado K, Cotelle V, Li G, Bellati J, Tang N, Tournaire-Roux C, Martiniere A, Santoni V, Maurel C. 2019. Oscillating Aquaporin Phosphorylation and 14-3-3 Proteins Mediate the Circadian Regulation of Leaf Hydraulics. Plant Cell 31: 417–429. Google Scholar: <u>Author Only Title Only Author and Title</u>

Rajakumara E, Law JA, Simanshu DK, Voigt P, Johnson LM, Reinberg D, Patel DJ, Jacobsen SE. 2011. A dual flip-out mechanism for 5mC recognition by the Arabidopsis SUVH5 SRA domain and its impact on DNA methylation and H3K9 dimethylation in vivo. Genes Dev 25: 137–152. Google Scholar: <u>Author Only Title Only Author and Title</u>

Rodriguez-Sanz H, Moreno-Romero J, Solis MT, Kohler C, Risueno MC, Testillano PS. 2014. Changes in histone methylation and acetylation during microspore reprogramming to embryogenesis occur concomitantly with BnHKMT and BnHAT expression and are associated with cell totipotency, proliferation, and differentiation in Brassica napus. Cytogenet Genome Res 143: 209–218. Google Scholar: <u>Author Only Title Only Author and Title</u>

Saze H, Shiraishi A, Miura A, Kakutani T. 2008. Control of genic DNA methylation by a jmjC domain-containing protein in Arabidopsis thaliana. Science 319: 462–465. Google Scholar: <u>Author Only Title Only Author and Title</u>

Scofield S, Murray JAH. 2006. KNOX gene function in plant stem cell niches. Plant Molecular Biology 60: 929–946. Google Scholar: <u>Author Only Title Only Author and Title</u>

Sinha NR, Williams RE, Hake S. 1993. Overexpression of the maize homeo box gene, KNOTTED-1, causes a switch from determinate to indeterminate cell fates. Genes Dev 7: 787–795. Google Scholar: <u>Author Only Title Only Author and Title</u>

Stroud H, Do T, Du J, Zhong X, Feng S, Johnson L, Patel DJ, Jacobsen SE. 2014. Non-CG methylation patterns shape the epigenetic landscape in Arabidopsis. Nature Structural & Molecular Biology 21: 64–72. Google Scholar: <u>Author Only Title Only Author and Title</u>

Thorstensen T, Fischer A, Sandvik SV, Johnsen SS, Grini PE, Reuter G, Aalen RB. 2006. The Arabidopsis SUVR4 protein is a nucleolar histone methyltransferase with preference for monomethylated H3K9. Nucleic Acids Res 34: 5461–5470. Google Scholar: <u>Author Only Title Only Author and Title</u>

Timmermans MCP, Hudson A, Becraft PW, Nelson T. 1999. ROUGH SHEATH2: A Myb protein that represses knox homeobox genes in maize lateral organ primordia. Science 284: 151–153. Google Scholar: <u>Author Only Title Only Author and Title</u>

Tran RK, Zilberman D, de Bustos C, Ditt RF, Henikoff JG, Lindroth AM, Delrow J, Boyle T, Kwong S, Bryson TD et al. 2005. Chromatin and siRNA pathways cooperate to maintain DNA methylation of small transposable elements in Arabidopsis. Genome Biol 6: R90. Google Scholar: <u>Author Only Title Only Author and Title</u>

Tsiantis M, Schneeberger R, Golz JF, Freeling M, Langdale JA. 1999. The maize rough sheath2 gene and leaf development programs in monocot and dicot plants. Science 284: 154–156. Google Scholar: <u>Author Only Title Only Author and Title</u>

Veiseth SV, Rahman MA, Yap KL, Fischer A, Egge-Jacobsen W, Reuter G, Zhou MM, Aalen RB, Thorstensen T. 2011. The SUVR4 histone lysine methyltransferase binds ubiquitin and converts H3K9me1 to H3K9me3 on transposon chromatin in Arabidopsis. PLoS Genet 7: e1001325. Google Scholar: <u>Author Only Title Only Author and Title</u>

Venglat SP, Dumonceaux T, Rozwadowski K, Parnell L, Babic V, Keller W, Martienssen R, Selvaraj G, Datla R. 2002. The homeobox gene BREVIPEDICELLUS is a key regulator of inflorescence architecture in Arabidopsis. Proc Natl Acad Sci U S A 99: 4730–4735. Google Scholar: <u>Author Only Title Only Author and Title</u>

Vial-Pradel S, Keta S, Nomoto M, Luo L, Takahashi H, Suzuki M, Yokoyama Y, Sasabe M, Kojima S, Tada Y et al. 2018. Arabidopsis Zinc-Finger-Like Protein ASYMMETRIC LEAVES2 (AS2) and Two Nucleolar Proteins Maintain Gene Body DNA Methylation in the Leaf Polarity Gene ETTIN (ARF3). Plant Cell Physiol 59: 1385–1397. Google Scholar: <u>Author Only Title Only Author and Title</u>

Vollbrecht E, Reiser L, Hake S. 2000. Shoot meristem size is dependent on inbred background and presence of the maize homeobox gene, knotted1. Development 127: 3161–3172. Google Scholar: <u>Author Only Title Only Author and Title</u>

Wang S, Yamaguchi M, Grienenberger E, Martone PT, Samuels AL, Mansfield SD. 2020. The Class II KNOX genes KNAT3 and KNAT7 work cooperatively to influence deposition of secondary cell walls that provide mechanical support to Arabidopsis stems. Plant J 101: 293–309. Google Scholar: <u>Author Only Title Only Author and Title</u>

Wang X, Liu S, Tian H, Wang S, Chen JG. 2015. The Small Ethylene Response Factor ERF96 is Involved in the Regulation of the Abscisic Acid Response in Arabidopsis. Front Plant Sci 6: 1064. Google Scholar: <u>Author Only Title Only Author and Title</u>

Xu L, Jiang H. 2020. Writing and reading histone H3 lysine 9 methylation in Arabidopsis. Front Plant Sci 11: 452. Google Scholar: <u>Author Only Title Only Author and Title</u>

Yang J, Yuan L, Yen MR, Zheng F, Ji R, Peng T, Gu D, Yang S, Cui Y, Chen PY et al. 2020. SWI3B and HDA6 interact and are required for transposon silencing in Arabidopsis. Plant J 102: 809–822. Google Scholar: <u>Author Only Title Only Author and Title</u>

You Y, Sawikowska A, Neumann M, Pose D, Capovilla G, Langenecker T, Neher RA, Krajewski P, Schmid M. 2017. Temporal dynamics of gene expression and histone marks at the Arabidopsis shoot meristem during flowering. Nat Commun 8: 15120. Google Scholar: <u>Author Only Title Only Author and Title</u>

Yu C-W, Liu X, Luo M, Chen C, Lin X, Tian G, Lu Q, Cui Y, Wu K. 2011. HISTONE DEACETYLASE6 interacts with FLOWERING LOCUS D and regulates flowering in Arabidopsis. Plant Physiology 156: 173–184. Google Scholar: <u>Author Only Title Only Author and Title</u>

Yu C-W, Tai R, Wang S-C, Yang P, Luo M, Yang S, Cheng K, Wang W-C, Cheng Y-S, Wu K. 2017. HISTONE DEACETYLASE6 acts in concert with histone methyltransferases SUVH4, SUVH5, and SUVH6 to regulate transposon silencing. Plant Cell 29: 1970–1983. Google Scholar: <u>Author Only Title Only Author and Title</u>

Yu G, Wang LG, He QY. 2015. ChIPseeker: an R/Bioconductor package for ChIP peak annotation, comparison and visualization. Bioinformatics 31: 2382–2383. Google Scholar: <u>Author Only Title Only Author and Title</u>

Zemach A, Kim MY, Hsieh P-H, Coleman-Derr D, Eshed-Williams L, Thao K, Harmer SL, Zilberman D. 2013. The Arabidopsis nucleosome remodeler DDM1 allows DNA methyltransferases to access H1-containing heterochromatin. Cell 153: 193–205. Google Scholar: <u>Author Only Title Only Author and Title</u>

Zhao L, Li Y, Xie Q, Wu Y. 2017. Loss of CDKC;2 increases both cell division and drought tolerance in Arabidopsis thaliana. Plant J 91: 816–828. Google Scholar: <u>Author Only Title Only Author and Title</u>

Zheng J, Chen F, Wang Z, Cao H, Li X, Deng X, Soppe WJJ, Li Y, Liu Y. 2012. A novel role for histone methyltransferase KYP/SUVH4 in the control of Arabidopsis primary seed dormancy. New Phytol 193: 605–616. Google Scholar: <u>Author Only Title Only Author and Title</u>

